# Dynamic surveillance of mosquitoes and their viromes in Wuhan during 2020

**DOI:** 10.1101/2021.07.17.452774

**Authors:** Nanjie Ren, Shunlong Wang, Chenyan Shi, Ping Yu, Lu Zhao, Doudou Huang, Haixia Ma, Shuqi Xiao, Fei Wang, Zhiming Yuan, Han Xia

## Abstract

Mosquitoes are medically important arthropod vectors and harbor a great variety of viruses. The population density, species and virome of mosquitoes varies according to geography and climate. To investigate the dynamic changes in the species composition and diversity of mosquitoes and their viromes in Wuhan, China, a total of 2,345 adult mosquitoes collected from different habitats including an urban residential area, two hospitals, a scenic area, and a pig farm in a rural region from April to October 2020 were subjected to morphological identification, RT-qPCR and metagenomic sequencing. The results indicated that the dominant presence of *Culex* mosquitoes was observed in both urban regions (90.32%, 1538/1703) and the pig farm (54.98%, 353/642). Viromes of *Culex* showed dynamic changes during the collection time. Several viruses, such as Culex flavivirus, Alphamesonivirus 1, Hubei mosquito virus 2 and Hubei mosquito virus 4, had seasonal changes and unimodal increases or declines. Other viruses, such as Wuhan mosquito virus 6, Hubei virga-like virus 2 and Zhejiang mosquito virus 3, were stable in all collected *Culex* and should be potential members of “core viromes”. This study improves the understanding of the dynamic composition of mosquitoes and the viromes they carry and provides useful information for informing mosquito control and mosquito-borne disease prevention strategies.

## Introduction

Mosquitoes, as blood-sucking arthropods, are widely distributed worldwide. Many of them cause severe threats to public health. Mosquitoes are vital competent vectors for several mosquito-borne viruses (MBVs), such as dengue virus and chikungunya virus, from which millions of people suffer every year [1][2]. In addition to MBVs that can infect both vertebrates and invertebrates, various mosquito-specific viruses (MSVs) have been identified, with possible applications as biological control agents against mosquito-borne diseases, diagnostic therapies and novel vaccine platforms [3]. With rapidly developing NGS (next-generation sequencing) technology, viral metagenomics has been used to detect viral diversity and abundance, predict disease outbreaks and identify novel viruses in uncultured mosquito samples [4]. Moreover, viral metagenomic surveillance systems have been established to monitor the virome derived from mosquito hosts [4].

Hubei is a province located in central China with a suitable environment for mosquito breeding. During the last decade, hundreds of Japanese encephalitis cases have been reported in Hubei [5]. In August 2019, the first local outbreak with a total of 50 cases of dengue fever was reported in Huangzhou, Hubei Province [6]. In addition, two genotypes of Banna virus (BAV) have been isolated from mosquitoes captured in Hubei; this virus is a member of the *Seadornavirus* genus and may be pathogenic to humans or animals [7]. Through metagenomic virome analysis, it was found that the viromes of mosquitoes from Hubei Province are mainly composed of MBVs, compared to the viromes of mosquitoes from Yunnan Province consisting of MSVs, suggesting that the composition of the virome is highly variable by geographic region [8]. For MSVs, Quang Binh virus, Culex flavivirus and Yichang virus (YCV), which have the ability to reduce the replication of DENV-2 when coinfected in mosquitoes, were also reported to be present in the mosquitoes from Hubei Province [9,10].

Wuhan is the capital city of Hubei Province, with an annual mean temperature of 15.8°C to 17.5°C and annual precipitation of 1,150 to 1,450 mm of rain, which makes this area suitable for mosquito breeding. During the COVID-19 epidemic, in early 2020, a large-scale disinfection procedure was applied within the environment of the whole city [11]. Therefore, compared to previous years, changes in the species or density of mosquito populations may occur in Wuhan. In addition, there is still limited information about the viral diversity and abundance of mosquitoes in Wuhan. In this study, mosquitoes were collected monthly from the field in Wuhan from April to October 2020. Dynamic surveillance of the mosquitoes and their virome and the detection of BAV and Japanese encephalitis virus (JEV) in mosquitoes were conducted.

## Methods and Materials

### Sampling sites

The sampling sites were selected in five representative regions of Wuhan, and the sampling times were from late April to late October 2020 (SFig. 1) as described previously [12]. The sites contain two hospitals, First People’s Hospital of Jiangxia District (30° 22′ 23″ N, 114° 18′ 56″ E) and Huoshenshan Hospital (30° 31′ 42″ N, 114° 4′ 50″ E); one urban residential area, Huanan seafood market (30° 37′ 7″ N, 114° 15′ 25″ E); one scenic area, East Lake (30° 37′ 6″ N, 114° 15′ 27″ E); and one pig farm in a rural region, Huangpi Pig Farm (30° 52′ 52″ N, 114° 22′ 30″ E).

**Fig. 1.**
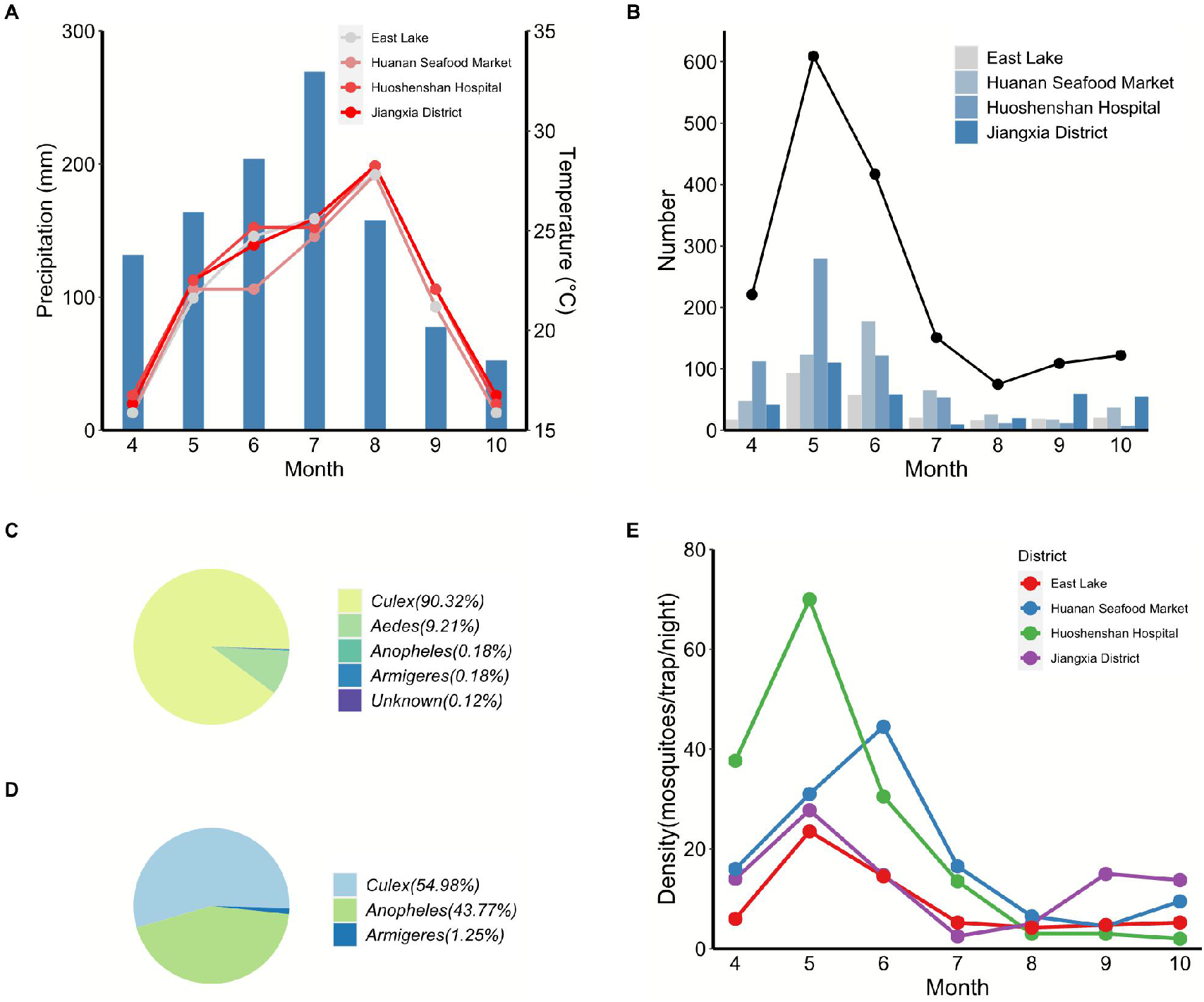
The species and amounts of mosquitoes collected in urban districts and Huangpi pig farm in Wuhan from April to October during 2020. (A) The histograms and lines show the precipitation and average temperature of each month, respectively. (B) The mosquitoes collected in urban districts per month. The line shows the total mosquitoes per month. (C) The proportion of mosquito genera in urban districts. (D) The proportion of mosquito genera in the Huangpi pig farm. (E) The abundance of mosquitoes at different collection times.

### Mosquito collection and sample preparation

Mosquitoes were collected using light traps with an attractant (Maxttrac, China) set on the shore ponds or in the bushes. Mosquitos were collected monthly from April to October, except in the pig farm, which was only trapped in May. The traps were set from 19:00 to 22:00 pm for 3-7 days every month. Then, the mosquitoes were separated, identified, and stored at -80°C.

The mosquitoes were assigned to different pools based on their respective collection site, month, species and sex. Each mosquito pool (20 mosquitoes/pool) was triturated by the cryogenic grinding method using a High-Speed Low-Temperature Tissue Grinding Machine (Servicebio, China) running two 30-second cycles at 50 Hz. After sufficient grinding, 600 μL of Roswell Park Memorial Institute (RPMI) medium was added for homogenization [13]. Mosquito macerates were clarified by centrifugation at 10,000 × g (4°C for 30 min) to remove cell debris and bacteria. Supernatants were stored at -80°C until further use.

### RNA extraction and RT-qPCR

RNA was extracted from 200 μL of homogenized supernatant sample using Direct-zol RNA MiniPrep (Zymo Research, USA) following the manufacturer’s instructions. The quantitative real-time reverse transcription PCR (RT-qPCR) mixtures for the detection of viral RNA were made by a Luna® Universal Probe One-Step RT-qPCR Kit (New England Biolabs, USA) in accordance with the manufacturer’s instructions and then placed in a thermocycler (BIO-RAD CFX96™ Real-Time System, USA). Specific RT-qPCR for BAV and JEV was performed using the reported primers and probes (STable 1)[7][14][15]. All oligoprimer DNAs were synthesized by TSINGKE (Wuhan Branch, China). Twenty-microliter reaction mixtures containing 5 μL of viral RNA and 0.8 μL of each primer were incubated at 55°C for 10 min and 95°C for 1 min followed by 40 cycles of 95°C for 10 s and 55°C for 30 s.

**Table 1.** Dominant mosquitoes in Wuhan based on metagenomic sequencing.

| Sampling site | Sample name | Mosquito genus | Month of collection | Number of mosquitoes for sequencing | Host-genome-removed reads (Viral proportion) | Number of nonredundant contigs aligned to eukaryotic viruses |
| --- | --- | --- | --- | --- | --- | --- |
| Urban district | whct-C_04 | *Culex* | April | 220 | 4,765,707 (5.90%) | 108 |
|  | whct-C_05 | *Culex* | May | 600 | 5,115,257 (19.26%) | 176 |
|  | whct-C_06 | *Culex* | June | 339 | 8,763,700 (11.46%) | 211 |
|  | whct-C_07 | *Culex* | July | 109 | 4,180,592 (12.96%) | 84 |
|  | whct-C_08 | *Culex* | August | 31 | 11,091,085 (3.50%) | 45 |
|  | whct-C_09 | *Culex* | September | 97 | 3,614,845 (10.60%) | 63 |
|  | whct-C_10 | *Culex* | October | 111 | 2,988,426 (6.02%) | 84 |
| Huangpi pig farm | hpzc-A_05 | *Anopheles* | - | 281 | 1,677,474 (20.14%) | 137 |
|  | hpzc-C_05 | *Culex* |  | 353 | 11,786,068 (3.37%) | 281 |

### Metagenomic sequencing

The dominant mosquito genera in urban districts and pig farms were selected for sequencing. Briefly, the RNA extracted from each mosquito pool was remixed to form new pools via different strategies. For urban pools, 10 μL of RNA extracted from pools from all the sampling areas in the same month was mixed into monthly pools. For the Huangpi pig farm pools, 10 μL of RNA extracted from pools from the same mosquito genus was mixed into *Anopheles* or *Culex* pools. The purity and integrity of the RNA pools were checked by the NanoPhotometer^®^ spectrophotometer (IMPLEN, CA, USA) and the RNA Nano 6000 Assay Kit and Bioanalyzer2100 system (Agilent Technologies, CA, USA). One microgram of RNA from each pool was used for library preparation with the NEBNext® UltraTMRNA Library Prep Kit for Illumina^®^ (NEB, USA) according to the manufacturer’s instructions. Then, the libraries were clustered on a cBot Cluster Generation System using a TruSeq PE Cluster Kit v3-cBot-HS (Illumina) and sequenced using an Illumina NovaSeq 6000 System.

### Downstream bioinformatics analysis

The paired-end reads from NGS were processed by Trim-galore [16] to trim adapters and low-quality bases, and the reads of the host genome (*Anopheles* or *Culex*) were discarded using bowtie2 [17] and bedtools [18]. The remaining reads were assembled into contigs by Trinityrnaseq [19]. The contigs longer than 500 bp were filtered and dereplicated (nucleotide identity > 95% and coverage rate > 80%) by CD-HIT [20] and aligned against the nonredundant protein database from NCBI (updated in March 2021) for taxonomic classification using Diamond blastX [21]. The BLASTX result was processed by the LCA algorithm (weighted LCA percent = 75%, e-value = 1 × 10^−5^, min Support = 1) with MEGAN [22]. Taxonomic classification was mainly conducted at the family level. The trimmed reads were aligned to the contigs of each sample by checkm [23] to calculate the abundance of eukaryotic viral contigs. Data visualization was conducted by ComplexHeatmap [24] and the ggplot2 [25] package in R [26]. A viral species that contained more than 500 reads was regarded to be present in a pool.

## Result

### Mosquito composition

The temperature and precipitation from April to October in Wuhan during 2020 (Fig. 1A) were recorded, and it was found that the rainfall increased in June and July, above the 10-year average data, but the average temperature was lower than that in previous years [27].

From April to October, a total of 2,345 adult mosquitoes were collected, with 1,703 from urban areas and 642 from the rural pig farm. The seasonal fluctuation of mosquito density in urban areas revealed a bimodal pattern, with the first peak occurring in May or June and the second significantly lower peak occurring in September or October (Fig. 1B and E), which was similar to previous data [28].

Four genera of mosquitoes were classified: *Culex, Aedes, Anopheles* and *Armigeres* (Fig. 1C-D). In these four genera, the predominant species in both urban and pig farms was *Culex*, which was consistent with existing reports [28]. In urban areas, *Culex* mosquitoes occupied a large proportion (90.32%, 1538/1703), followed by *Aedes* (9.21%, 157/1703). Both *Anopheles* (0.18%, 3/1703) and *Armigeres* (0.18%, 3/1703) were caught in small amounts. However, on pig farms, the main mosquito species were *Culex* (54.98%, 353/642) and *Anopheles* (43.77%, 281/1703). *Armigeres* (1.25%, 8/1703) was much less collected, while no *Aedes* was caught.

### Virus detection by RT-qPCR

The 2,345 collected mosquitoes were separated into 123 pools, with 90 from urban areas and 33 from the pig farm. From 90 urban pools, BAV was detected in 18, with Ct values ranging from 32 to 38, and the sequencing results confirmed that 14 out of 18 samples were BAV positive. The BAV-positive samples were mainly distributed in residential areas from May to July. For the pig farm, only one sample collected in May was BAV positive (Ct = 32.71) and confirmed by sequencing. However, none of the mosquito pools were JEV positive.

### Virome profile of mosquitoes collected in Wuhan

The seven *Culex* monthly pools (whct-C_04 to whct-C_10) from urban areas and one *Anopheles* (hpzc-A_05) and one *Culex* pool (hpzc-C_05) from the Huangpi pig farm were processed for metagenomic sequencing. All nine pools contained 53,983,154 host-genome-removed reads in total (Table 1). A total of 2,663,238 de novo assembled contigs were added. The contigs were clustered and dereplicated (longer than 500 bp, nucleotide identity > 95% and coverage > 80%) into 413,645 nonredundant contigs. BLASTX results showed that 1,189 and 412,456 contigs were aligned to eukaryotic viruses and other organisms, respectively.

The viromes of the collected mosquitoes were classified into 20 viral families and a group of unclassified viruses (Fig. 2A). Invertebrate viruses were dominant in the viromes, and the hosts of 11 viruses remain unknown (Fig. 2B). Only Hubei chryso-like virus 1 (HCLV1) has a vertebrate host.

**Fig. 2.**
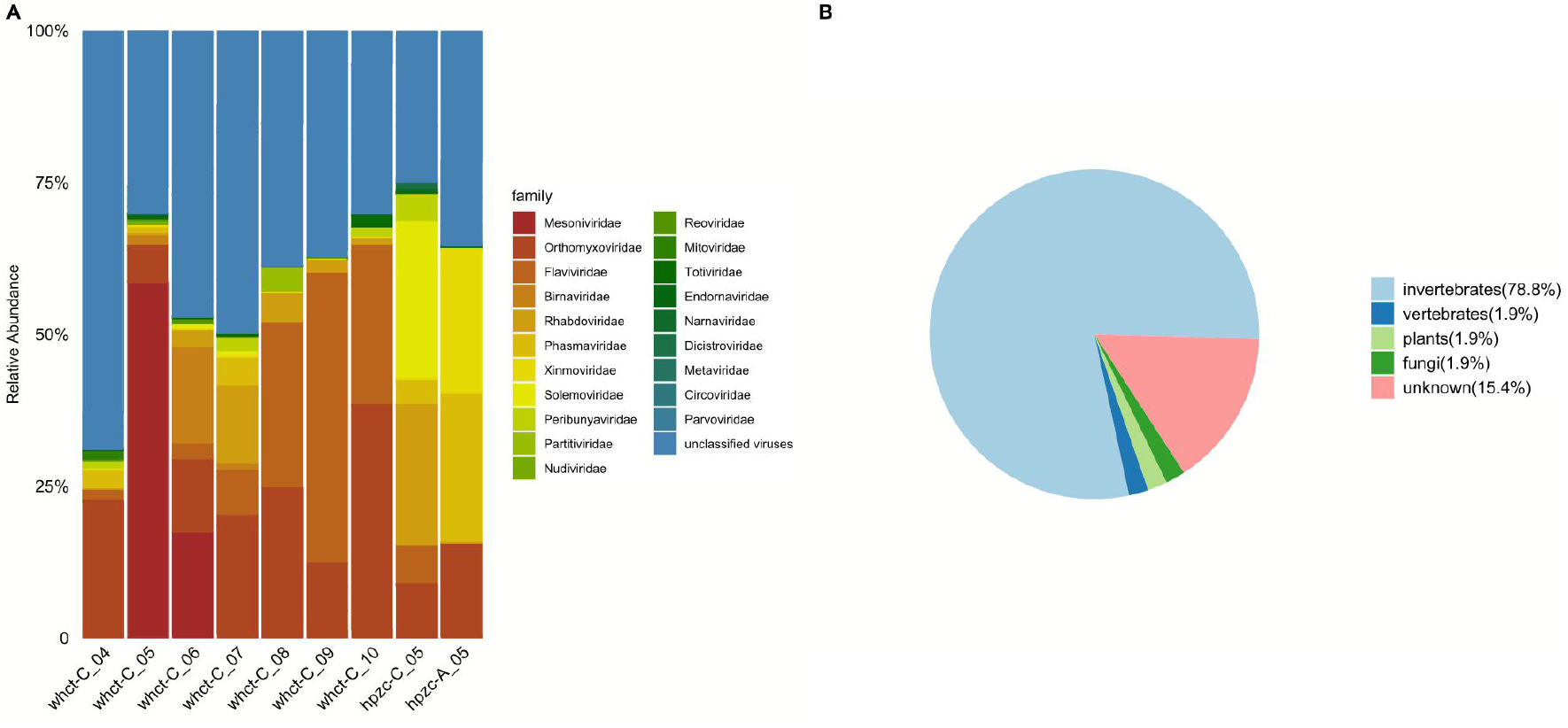
Families and hosts of viruses in the viromes of mosquitoes in different pools. (A) The map shows the relative abundance of viral families in each pool. (B) The composition of the hosts of all 52 viral species.

The abundance of each viral species in which at least one pool contained more than 500 reads is shown in the heatmap (Fig. 3). The viromes contained 52 viral species in total. Thirty of them could be classified into 20 viral families, and the other 22 viruses lacked accurate taxonomic classification. Unclassified viruses composed a large proportion of the viromes in all pools.

**Fig. 3.**
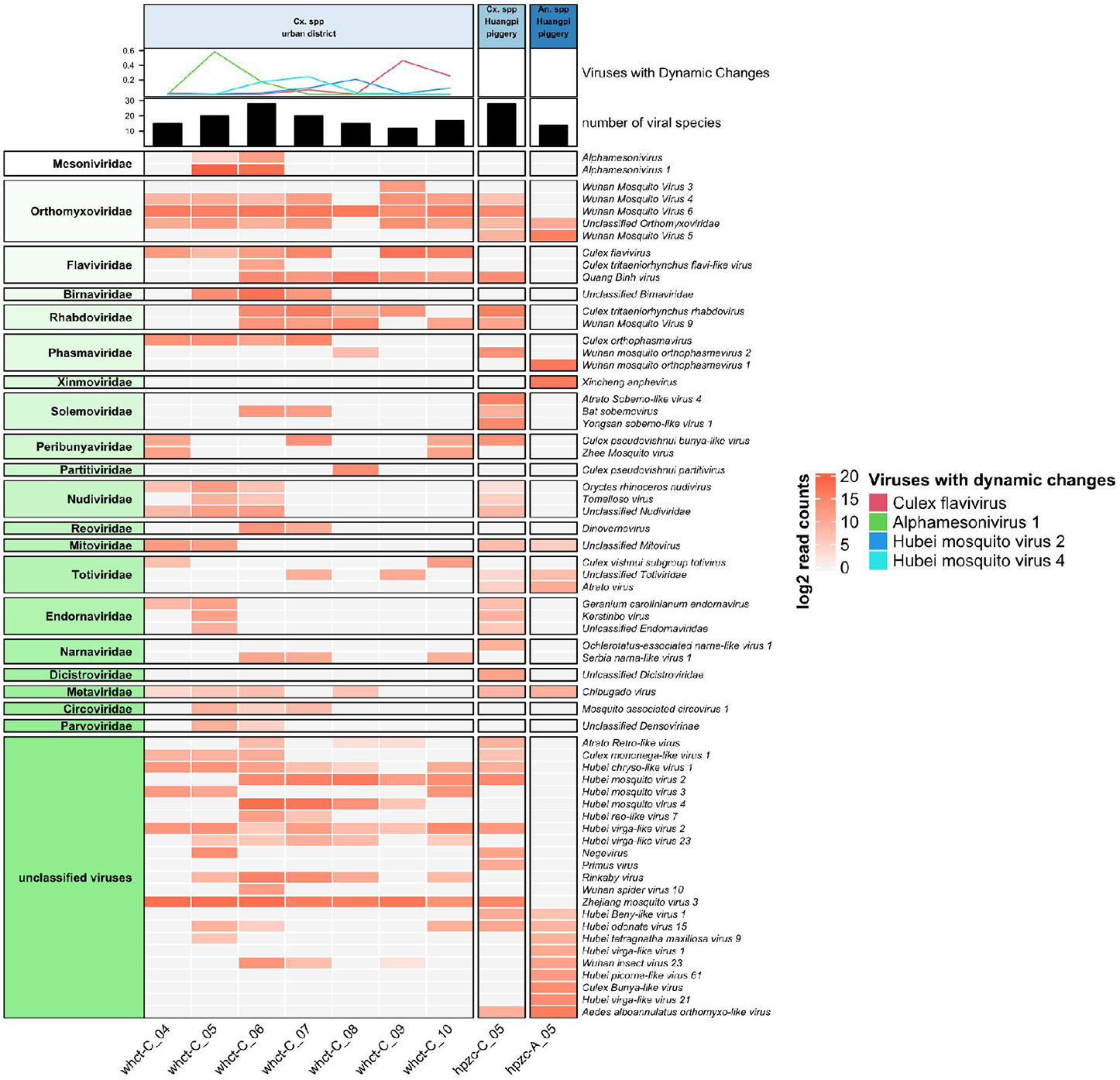
The abundance of viruses in each pool. Viral species contained more than 500 reads in at least one pool. The bar plot shows the number of viral species in each sample. The line plot at the top shows four viruses that had dynamic changes at different collection times.

Eighteen viral families were present in viromes of *Culex* mosquito pools from urban districts. Viral diversity in urban districts was altered by month and showed two peaks. The highest viral diversity occurred in the pool whct-C_06 (Fig. 3A). Several viruses, such as Culex flavivirus and Zhejiang mosquito virus 3, had significant changes in relative abundance in these months. The family *Orthomyxoviridae* had the most reads in the pools from urban districts and contained four members (Wuhan mosquito virus 3, Wuhan mosquito virus 4, Wuhan mosquito virus 5 and Wuhan mosquito virus 6) that were recently identified in Hubei Province (Fig. 3). Culex flavivirus and Quang Binh virus, both MSVs, were present in the pools. The *Culex* pool of the Huangpi pig farm, which contained 13 viral families and 4 unique viruses (Atrato Sobemo-like virus 4, Yongsan sobemo-like virus 1, Primus virus and Ochlerotatus-associated narna-like virus), had viromes similar to those of urban districts. In all *Culex* pools, Wuhan Mosquito virus 6, Hubei virga-like virus 2 and Zhejiang mosquito virus 3 could be found. The differences in viral composition between the urban and rural areas was mainly found for viral families such as *Parvoviridae* and *Circoviridae*.

The viromes in *Anopheles* mosquitoes were significantly lower than those in *Culex* mosquitoes. The *Anopheles* pool had six unique viruses, and the most abundant virus was Xincheng anphevirus. Nine viral species could be found in the two mosquito species from the Huangpi pig farm, four of which (Wuhan mosquito virus 5, Atrato virus, Hubei Beny-like virus 1 and *Aedes alboannulatus* orthomyxo-like virus) were absent in *Culex* mosquitoes in urban districts.

## Discussion

MBVs are pervasive public health problems on a global scale, and effective surveillance programs for both vectors and pathogens are needed. In this study, we completed 7 months of mosquito monitoring in Wuhan during 2020 to determine the dynamic changes in mosquito populations and their viromes.

Between-site habitat differences, climates and longer seasonal fluctuations can influence the number of mosquitoes [29]. Although efforts were made to sample throughout the whole year, the heavy rainfall from June to September did reduce the sampling quantity of mosquitos in Wuhan. The highest abundances of the mosquito catch in urban areas were in May, but that number plummeted in August (Fig. 1). Apart from the highest precipitation in July and the highest temperature in August, the low abundances of mosquito catches may also be due to the disinfection strategies carried out by the government in July for students returning to school and public entertainment reopening.

Compared to the trapped mosquitoes from 2017 to 2019 in Wuhan, which can be classified into 8 species under 5 genera and 1 family [28], the diversity of mosquito species was lower in this study (Fig. 1). This result could be attributed to the chemical blend of the attractant not resulting in trap catches as high as those elicited by human odor [30]. Furthermore, the morphological quality of mosquitoes deteriorated in large and soft nets used in this study, resulting in mosquitoes that could only be classified to the genus level here. Further investigations into diversity were prevented, which could be the other reason for the lower diversity of mosquito genera.

BAV belongs to the family of *Reoviridae*, RT-qPCR results showed that BAV existed in mosquitoes from both urban areas and rural pig farms in Wuhan. Although BAV contigs was not detected in the mosquito pools for NGS sequencing, there were many unclassified reads in *Reoviridae* in whct-C_06 and whct-C_07, which corresponded with qPCR detection for BAV, probably due to the different pooling strategies in the two methods.

Viral metagenomics has been applied in the analysis of viromes from mosquitoes in many studies [8,31,32]. By NGS, 3.37% -20.14% reads from each pool collected from urban districts and a pig farm in Wuhan were aligned to viruses (Table 1). Additionally, *Orthomyxoviridae* and *Flaviviridae* were prevalent in *Culex* viromes, in which invertebrate viruses were dominant. Previous studies performed in Hubei Province showed that the prevalent viral families in *Culex* mosquitoes are *Herpesviridae* and *Adenoviridae*, whose hosts are vertebrates [8].

The dynamic changes in the viromes of *Culex* from urban districts in Wuhan were monitored (Fig. 3). The viral diversity in the monthly pools followed a bimodal distribution, and the two peaks occurred in June and October. A previous study performed in the USA and Trinidad showed that several insect-specific viruses have evident seasonal activity [33]. The viral diversity in different months in Wuhan may also have seasonal changes. In our present study, a large proportion of viral families were absent in several pools, probably due to their low abundance in all pools. *Orthomyxoviridae* and *Flaviviridae* were the dominant viral families and stably existed in the monthly pools (Fig. 3). The relative abundance of *Orthomyxoviridae* had no obvious trend during all months but increased from May to August. Notably, four virus species (Culex flavivirus, Alphamesonivirus 1, Hubei mosquito virus 2 and Hubei mosquito virus 4) had significant changes in these months. Culex flavivirus showed a low level of abundance in April to July and had higher abundance in September to October. Alphamesonivirus 1, Hubei mosquito virus 2 and Hubei mosquito virus 4 show unimodal curves, and their peaks occur in May, August and June, respectively. Several of these viruses were highly abundant in a few months but disappeared in the other months, suggesting that they may have a strong correlation with the alternation of seasons and possibly related to the activity of the *Culex* population. Unfortunately, environmental factors were not sufficiently recorded, which limited further exploration of the correlation between mosquito viromes and many other factors, such as climate and host composition.

In the heatmap (Fig. 3), Wuhan mosquito virus 3, Wuhan mosquito virus 4, Wuhan mosquito virus 5 and Wuhan mosquito virus 6 belong to the genus *Quaranjavirus*, family *Orthomyxoviridae* [34]. Several members of this genus, such as Quaranfil virus and Johnston Atoll virus, can infect both vertebrate and invertebrate hosts and result in the death of newborn mice in the laboratory [35]. The other members of *Quaranjavirus* have not been properly characterized, and it should be confirmed whether they are potential pathogenic to humans and livestock. Quang Binh virus, Culex tritaeniorhynchus flavi-like virus (CtFLV) and Culex flavivirus, which belong to the family *Flaviviridae*, have variable distributions in many regions [36][37]. Quang Binh virus was first isolated in Vietnam and then chronologically reported in Yunnan Province and Hubei Province in China [8][38]. Recently, it was reported to be detected in *Culex* from northwestern China, suggesting that the virus may have an expanded geographical distribution [31]. Several arboviruses, such as CtFLV, identified from the local *Culex* in Japan, were closely related to those identified in Hubei Province [37]. In consideration of the frequent exchanges between China and Japan, mosquitoes can be transferred through human activities. It is worth noting whether viromes of mosquitoes interact between the two locations.

Viromes of mosquitoes in Wuhan reflected geographic and species differences. *Culex* species from urban districts and the pig farm had similar viral family compositions. Rhabdoviriuses were most prevalent in the pig farm, and several unclassified viruses only existed in one of the two locations, which was probably due to diverse mosquito hosts and environments in urban and rural areas [39]. Wuhan mosquito orthophasmavirus 1 (Family *Phasmaviridae*) and Xincheng anphevirus (Family *Xinmoviridae*) dominated in *Anopheles*; both are insect-specific viruses. *Anopheles* and *Culex* had significantly distinct viromes, which is similar to previous studies in Guadeloupe and Japan [32][37]. Wuhan mosquito virus 6, Hubei virga-like virus 2 and Zhejiang mosquito virus 3 were present in all the *Culex* pools (Fig. 3). Several studies have reported these viruses in mosquitoes from multiple geographical regions, suggesting that they are potential members of the “core viromes” of mosquitoes [32][40]. Widely distributed MSVs may play a vital role in the biocontrol of mosquitoes. In the unclassified virus group, HCLV1 has a vertebrate host, but the evidence is insufficient because HCLV1 was isolated from the feces of *Cairina moschata* (also known as Muscovy duck) in Australia [41]. The transmission of this virus is still unknown, and many more studies are needed to explore whether birds act as hosts since Muscovy ducks are widely bred in Hubei.

## Acknowledgements

This work was supported by the the Wuhan Science and Technology Plan Project (2018201261638501) and Open Foundation of Key Laboratory of Tropical Translational Medicine of Ministry of Education, Hainan Medical University (2021TTM010).

## Reference

1. Souza-Neto JA, Powell JR, Bonizzoni M: Aedes aegypti vector competence studies: A review. Infect Genet Evol. 2019; 67:191–209.

2. Arcà B, Colantoni A, Fiorillo C, et al.: MicroRNAs from saliva of anopheline mosquitoes mimic human endogenous miRNAs and may contribute to vector-host-pathogen interactions. Sci Rep. 2019; 9:.

3. Atoni E, Zhao L, Karungu S, et al.: The discovery and global distribution of novel mosquito-associated viruses in the last decade (2007-2017). Rev Med Virol. 2019; 29:.

4. Brinkmann A, Nitsche A, Kohl C: Viral metagenomics on blood-feeding arthropods as a tool for human disease surveillance. Int J Mol Sci. 2016; 17:.

5. Inspection H, Province H: Analysis of Laboratory Confirmed Case of Epidemic Encephalitis B in Hubei Province from 2011 to 2016. 2017; 14–16.

6. Dan-qin H, Li LIU, Qi C, et al.: Analysis of dengue epidemic and Aedes vector surveillance in Hubei province,China,2019. 2021; 32:38–40.

7. Xu L hong, Cao Y xi, He L fang, et al.: Detection of Banna virus-specific nucleic acid with TaqMan RT-PCR assay. Zhonghua Shi Yan He Lin Chuang Bing Du Xue Za Zhi. 2006; 20:47–51.

8. Atoni E, Wang Y, Karungu S, et al.: Metagenomic virome analysis of aed mosquitoes from Kenya and China. Viruses. 2018; 10:.

9. Wang Y, Xia H, Zhang B, Liu X, Yuan Z: Isolation and characterization of a novel mesonivirus from Culex mosquitoes in China. Virus Res. 2017; 240:130–139.

10. Atoni E, Zhao L, Hu C, et al.: A dataset of distribution and diversity of mosquito-associated viruses and their mosquito vectors in China. Sci Data. 2020; 7:.

11. Zhou P, Shi Z: SARS-CoV-2 spillover events. Science. 2021; 371:120–122.

12. Zhao L, Atoni E, Nyaruaba R, et al.: Environmental surveillance of SARS-CoV-2 RNA in wastewater systems and related environments in Wuhan: April to May of 2020. J Environ Sci. 2022; 112:115–120.

13. Wang J, Li H, He Y, et al.: Isolation of Tibet orbivirus from Culicoides and associated infections in livestock in Yunnan, China. Virol J. 2017; 14:105.

14. Pyke AT, Smith IL, Van Den Hurk AF, et al.: Detection of Australasian Flavivirus encephalitic viruses using rapid fluorogenic TaqMan RT-PCR assays. J Virol Methods. 2004; 117:161–167.

15. Pyke AT, Williams DT, Nisbet DJ, et al.: The appearance of a second genotype of Japanese encephalitis virus in the Australasian region. Am J Trop Med Hyg. 2001; 65:747–753.

16. Babraham Bioinformatics - Trim Galore! no date; .

17. Langmead B, Salzberg SL: Fast gapped-read alignment with Bowtie 2. Nat Methods. 2012; 9:357–359.

18. Quinlan AR: BEDTools: The Swiss-Army tool for genome feature analysis. Curr Protoc Bioinforma. 2014; 2014:11.12.1-11.12.34.

19. Haas BJ, Papanicolaou A, Yassour M, et al.: De novo transcript sequence reconstruction from RNA-seq using the Trinity platform for reference generation and analysis. Nat Protoc. 2013; 8:1494–1512.

20. Fu L, Niu B, Zhu Z, Wu S, Li W: CD-HIT: Accelerated for clustering the next-generation sequencing data. Bioinformatics. 2012; 28:3150–3152.

21. Buchfink B, Reuter K, Drost HG: Sensitive protein alignments at tree-of-life scale using DIAMOND. Nat Methods. 2021; 18:366–368.

22. Huson DH, Beier S, Flade I, et al.: MEGAN Community Edition - Interactive Exploration and Analysis of Large-Scale Microbiome Sequencing Data. PLoS Comput Biol. 2016; 12:1004957.

23. Parks DH, Imelfort M, Skennerton CT, Hugenholtz P, Tyson GW: CheckM: Assessing the quality of microbial genomes recovered from isolates, single cells, and metagenomes. Genome Res. 2015; 25:1043–1055.

24. Gu Z, Eils R, Schlesner M: Complex heatmaps reveal patterns and correlations in multidimensional genomic data. Bioinformatics. 2016; 32:2847–2849.

25. Wickham H: ggplot2: Elegant Graphics for Data Analysis. Springer-Verlag New York, 2016.

26. Team C: R: A Language and Environment for Statistical Computing. 2015.

27. National Greenhouse Data System: no date; .

28. Liu J, Mao W, Ding H, et al.: Composition and diversity of mosquito community in Wuhan from 2017 to 2019. J Cent China Norm Univ. 2021; 55:416–423.

29. Eastwood G, Sang RC, Lutomiah J, Tunge P, Weaver SC: Sylvatic mosquito diversity in kenya—considering enzootic ecology of arboviruses in an era of deforestation. Insects. 2020; 11:.

30. Jawara M, Awolola TS, Pinder M, et al.: Field testing of different chemical combinations as odour baits for trapping wild mosquitoes in the Gambia. PLoS One. 2011; 6:.

31. He X, Yin Q, Zhou L, et al.: Metagenomic sequencing reveals viral abundance and diversity in mosquitoes from the Shaanxi-Gansu-Ningxia region, China. PLoS Negl Trop Dis. 2021; 15:.

32. Shi C, Beller L, Deboutte W, et al.: Stable distinct core eukaryotic viromes in different mosquito species from Guadeloupe, using single mosquito viral metagenomics. Microbiome. 2019; 7:121.

33. Kim DY, Guzman H, Bueno R, et al.: Characterization of Culex Flavivirus (Flaviviridae) strains isolated from mosquitoes in the United States and Trinidad. Virology. 2009; 386:154–159.

34. Li CX, Shi M, Tian JH, et al.: Unprecedented genomic diversity of RNA viruses in arthropods reveals the ancestry of negative-sense RNA viruses. Elife. 2015; 2015:.

35. Presti RM, Zhao G, Beatty WL, et al.: Quaranfil, Johnston Atoll, and Lake Chad Viruses Are Novel Members of the Family Orthomyxoviridae . J Virol. 2009; 83:11599–11606.

36. Agboli E, Leggewie M, Altinli M, Schnettler E: Mosquito-specific viruses— transmission and interaction. Viruses. 2019; 11:.

37. Faizah AN, Kobayashi D, Isawa H, et al.: Deciphering the virome of culex vishnui subgroup mosquitoes, the major vectors of japanese encephalitis, in Japan. Viruses. 2020; 12:264.

38. Crabtree MB, Nga PT, Miller BR: Isolation and characterization of a new mosquito flavivirus, Quang Binh virus, from Vietnam. Arch Virol. 2009; 154:857–860.

39. Hui G, Huagang L, Lin W, et al.: The structure, spaciotemporal dynamics, and diversity of mosquito communities in Wuhan. Chinese J Appl Entomol. 2020; 57:955–962.

40. Shi C, Zhao L, Atoni E, et al.: Stability of the Virome in Lab- and Field-Collected Aedes albopictus Mosquitoes across Different Developmental Stages and Possible Core Viruses in the Publicly Available Virome Data of Aedes Mosquitoes . MSystems. 2020; 5:.

41. Vibin J, Chamings A, Collier F, Klaassen M, Nelson TM, Alexandersen S: Metagenomics detection and characterisation of viruses in faecal samples from Australian wild birds. Sci Rep. 2018; 8:.

