## Supplemental Table 1 for "Dynamic surveillance of mosquitoes and their viromes in Wuhan during 2020"

**STable 1. Primers and probes used for testing**

| Virus | Target | Forward primer | Reverse primer | Probe |
|---|---|---|---|---|
| BAV | VP10 | 5′-TCGGGCAACCATGCTTTCC-3′ | 5′-GTTGGTAGAGGGTGGTTGACATC-3′ | 5′-FAM-CGGGATCCTCTACTTTCCAGTCCCAAGA-3′-BHQ1 |
| JEV | NS5 | 5′-ATCTGGTGYGGYAGTCTCA-3′ | 5′-CGCGTAGATGTTCTCAGCCC-3′ | 5′-FAM-CGGAACGCGATCCAGGGCAA-3′-BHQ1 |
