## Supplementary figures and images for "Dynamic surveillance of mosquitoes and their viromes in Wuhan during 2020"

### Supplemental Fig 1

**SFig. 1 Sampling area in Wuhan.**

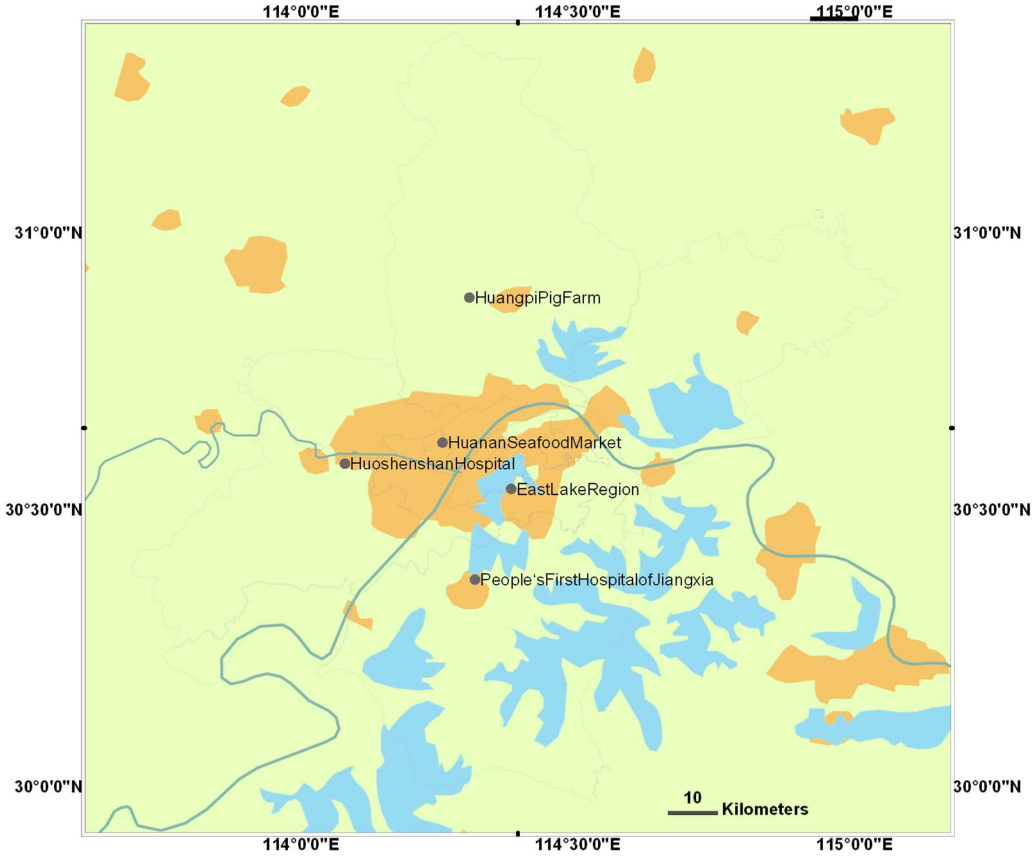
